# Lower cognitive set shifting ability is associated with stiffer balance recovery behavior and larger perturbation-evoked cortical responses in older adults

**DOI:** 10.1101/2021.07.16.452646

**Authors:** Aiden M. Payne, Jacqueline A. Palmer, J. Lucas McKay, Lena H. Ting

**Author notes:** **Correspondence:** L. H. Ting.

## Abstract

The mechanisms underlying associations between cognitive set shifting impairments and balance dysfunction are unclear. Cognitive set shifting refers to the ability to flexibly adjust behavior to changes in task rules or contexts, which could be involved in flexibly adjusting balance recovery behavior to different contexts, such as the direction the body is falling. Prior studies found associations between cognitive set shifting impairments and severe balance dysfunction in populations experiencing frequent falls. The objective of this study was to test whether cognitive set shifting ability is expressed in successful balance recovery behavior in older adults with high clinical balance ability (N=19, 71 ± 7 years, 6 female). We measured cognitive set shifting ability using the Trail Making Test and clinical balance ability using the miniBESTest. For most participants, cognitive set shifting performance (Trail Making Test B-A = 37 ± 20s) was faster than normative averages (46s for comparable age and education levels), and balance ability scores (miniBESTest = 25 ± 2 / 28) were above the threshold for fall risk (23 for people between 70-80 years). Reactive balance recovery in response to support-surface translations in anterior and posterior directions was assessed in terms of body motion, muscle activity, and brain activity. Across participants, lower cognitive set shifting ability was associated with smaller peak center of mass displacement during balance recovery, lower directional specificity of late phase balance-correcting muscle activity (i.e., greater antagonist muscle activity 200-300ms after perturbation onset), and larger cortical N1 responses (100-200ms). None of these measures were associated with clinical balance ability. Our results suggest that cognitive set shifting ability is expressed in balance recovery behavior even in the absence of profound clinical balance disability. Specifically, our results suggest that lower flexibility in cognitive task performance is associated with lower ability to incorporate the directional context into the cortically-mediated later phase of the motor response. The resulting antagonist activity and stiffer balance behavior may help explain associations between cognitive set shifting impairments and frequent falls.

## 2. INTRODUCTION

Cognitive impairment is associated with balance dysfunction, but it is unclear whether or how cognitive ability relates to balance recovery behavior in relatively high-functioning preclinical populations. Subtle cognitive impairments in executive function (Muir et al., 2012), attention, and memory are associated with clinical balance impairments (Tangen et al., 2014) and predict the first (Herman et al., 2010) and recurring falls in older adults (Gleason et al., 2009; Mirelman et al., 2012). However, it is unclear whether subtle differences in cognitive ability in the absence of clinically detectable balance dysfunction are associated with changes in balance control. Associations between cognitive function and balance control could provide mechanistic insight into findings that cognitive engagement in balance control increases with age (Rankin et al., 2000), fall history (Shumway-Cook et al., 1997), and fall risk (Lundin-Olsson et al., 1997). Here, we focus on individual differences in cognitive set shifting ability (i.e., the ability to flexibly adjust a behavior to changes in task rules or contexts), which have previously been associated with clinical balance dysfunction (Tangen et al., 2014), fall history (McKay et al., 2018), and fall risk (Herman et al., 2010). We investigate balance recovery behavior in terms of body motion, muscle activity, and brain activity evoked by a sudden balance disturbance, in contrast to prior studies that used clinical instruments and falls tracking, which are less applicable to preclinical populations. Identifying associations between cognitive ability and balance recovery behavior in preclinical populations could provide insight into underlying mechanisms for balance impairments that could serve as therapeutic targets for rehabilitation prior to occurrence of a fall.

Cognitive set shifting is an executive function that pertains to the ability to flexibly adjust behavior to changes in task rules or contexts, but its potential role in balance behavior is unclear. The Trail Making Test is a common pen and paper assessment of cognitive set shifting, consisting of two parts. In Part A, a participant must rapidly draw lines to connect dots in numerical order, relying on sustained attention, working memory, visuomotor search, and dexterity (Sanchez-Cubillo et al., 2009). In Part B, the task is altered to incorporate switching between numbers and letters (1-A-2-B-3-C…), thereby adding in a component of cognitive set shifting (Sanchez-Cubillo et al., 2009). Scoring the difference in time to complete Part B – Part A accounts for the overlapping motor and cognitive aspects, leaving a relatively pure measure of cognitive set shifting (Sanchez-Cubillo et al., 2009). However, the construct of set shifting inherently includes an increased working memory load to maintain and switch between two rule sets, as well as response selection and inhibition to select the response according to the current rule set while suppressing the response to the previous rule set (Koch et al., 2010). Because the Trail Making Test is so far removed from standing balance behavior, it is unclear why it has been repeatedly associated with advanced balance impairments (Herman et al., 2010; Tangen et al., 2014; McKay et al., 2018). However, if the neural mechanisms for cognitive set shifting assessed by the Trail Making Test are involved in balance control, then variation in cognitive set shifting ability should be expressed in successful balance recovery behavior before people begin experiencing frequent falls.

Similar to effective cognitive control, successful balance recovery behavior requires quick and flexible execution of a contextually appropriate behavior. Support-surface translational perturbations rapidly displace the base of support (i.e., the feet) relative to the body’s center of mass, requiring a rapid neural and mechanical reaction to prevent a fall. Effective balance recovery behavior involves directionally specific motor responses, with muscles showing preferential activation in response to perturbation directions in which they can generate torque to counteract center of mass displacement (Henry et al., 1998; Torres-Oviedo and Ting, 2007). This type of directional specificity is reduced in people with balance impairments (Lang et al., 2019), resulting in simultaneous agonist-antagonist cocontraction, which increases joint stiffness, but ultimately limits joint torques as the actions of the agonist and antagonist muscles partially resist one another (Damiano, 1993). Although cocontraction is common when learning new or complex motor skills and can be beneficial in some contexts (Damiano, 1993), its association to balance impairments suggests it is not an ideal strategy for balance recovery behavior (Lang et al., 2019). It has been suggested that cognitive flexibility, a broader construct containing cognitive set shifting, may be needed to quickly adjust behavior to unpredictable demands, including the use of feedback from the body or environment to appropriately react to a sudden displacement of the body’s center of mass (Pieruccini-Faria et al., 2019). Here, we test whether cognitive set shifting ability is associated with the ability to modulate muscle activity between balance perturbations that displace the body’s center of mass in opposite directions.

Testing different phases of the motor response for associations to cognitive ability could provide insight into different mechanisms by which cognitive function may overlap with balance recovery behavior. A sudden balance perturbation evokes a relatively stereotyped brainstem-mediated balance-correcting motor response at ∼100 ms via integrated sensory inputs reflecting the task-level goal of upright posture, and not the local stretch of individual muscles (Nashner, 1979; Horak and Nashner, 1986; Dietz et al., 1987; Safavynia and Ting, 2013b). While this early response is subcortically-mediated, it can be influenced by pre-perturbation cognitive state, including arousal (Carpenter et al., 2004), expectations (Horak et al., 1989), and intentions (McIlroy and Maki, 1993; Burleigh et al., 1994; Burleigh and Horak, 1996; Weerdesteyn et al., 2008) in ways that may depend on descending cortical influence in anticipation of an upcoming balance disturbance. More variable motor responses occur at longer latencies (>150 ms) that can incorporate cortically-mediated motor responses to the balance disturbance (Jacobs and Horak, 2007a). Cognitive dual task interference is limited to this later phase of the motor response, suggesting only the later phase depends on online cognitive processing (Rankin et al., 2000). If cognitive set shifting is associated with directional specificity in the early phase of the response, this would implicate cognitive set shifting in the maintenance of “central set,” which refers to the ability of the central nervous system to preselect the gain of stimulus-evoked behaviors in consideration of arousal, expectations, and intentions (Prochazka, 1989). If cognitive set shifting is associated with directional specificity only in the later phase, this would implicate cognitive set shifting in cortically-mediated reactions to the balance perturbation, which can incorporate incoming sensory information into decisions about how to react.

Balance perturbations also evoke a cortical response that is associated with balance ability and cognitive processing, but it is unknown whether this cortical response reflects individual differences in cognitive ability. A cortical response, termed the “N1” for the first negative peak in the evoked electroencephalography signal, occurs in the supplementary motor area ∼150 ms after a balance disturbance (Marlin et al., 2014; Mierau et al., 2015). We have previously suggested that the cortical N1 may reflect compensatory cortical engagement in balance recovery because it is enhanced in young adults with lower balance ability (Payne and Ting, 2020a) and on trials in which compensatory steps are taken (Payne and Ting, 2020c). The cortical N1 is also influenced by cognitive processes including attention (Quant et al., 2004b; Little and Woollacott, 2015), perceived threat (Adkin et al., 2008; Mochizuki et al., 2010), and predictability (Adkin et al., 2006; Adkin et al., 2008; Mochizuki et al., 2008; Mochizuki et al., 2010) and may therefore reflect cognitive-motor interactions. The possibility that the N1 reflects cognitive-motor interactions is further supported by its localization to the supplementary motor area (Marlin et al., 2014; Mierau et al., 2015), which is thought to mediate interactions between cognitive and motor processes by mediating interactions between neighboring prefrontal and motor cortical areas (Goldberg, 1985). Although investigations of the cortical N1 in older populations have been limited (Duckrow et al., 1999; Ozdemir et al., 2018), the N1 may be ideally suited for investigating relationships between cognitive and motor impairments with aging. Here, we test whether the cortical N1 is associated with individual differences in cognitive set shifting ability.

We investigated whether individual differences in cognitive set shifting ability were associated with perturbation-evoked balance recovery behavior and cortical activity in an older population with relatively high balance function to gain insight into possible mechanisms linking balance and cognitive function. We assessed clinical balance ability with the mini Balance Evaluation Systems Test (miniBESTest) (Magnani et al., 2020) and cognitive set shifting ability with the Trail Making Test (Tombaugh, 2004; Sanchez-Cubillo et al., 2009). We tested these ability measures for association with perturbation-evoked balance recovery behavior, including whole body stiffness, directional specificity of ankle muscle activity in early and late phases of the motor response, and the evoked cortical N1 response. We found that individuals with lower cognitive set shifting ability had stiffer behavior, less directional specificity of muscle activity, and larger cortical responses, revealing aspects of behavior that may share neural mechanisms involved in cognitive set shifting behavior.

## 3. METHODS

### 3.1 Participants

Nineteen older adults (age 71±6, 6 female) participated in this study. Written consent was obtained from all participants after a detailed explanation of the protocol according to procedures approved by the Emory University Institutional Review Board.

Participants were recruited from Emory University and the surrounding community. Adults over 55 years of age were screened for the following inclusion criteria: vision can be corrected to 20/40 or better with corrective lenses, no history of stroke or other neurologic condition, no musculoskeletal conditions or procedures that cause pain or limit mobility of the legs, ability to stand unassisted for at least 15 minutes, and cognitive ability to provide informed consent. Potential participants were excluded for prior experience on the perturbation platform. Study data were collected and managed using a Research Electronic Data Capture (REDCap) database hosted at Emory University (Harris et al., 2009; Harris et al., 2019).

### 3.2 Balance ability

The miniBESTest (www.bestest.us) was used as a measure of balance ability (Magnani et al., 2020) which assesses anticipatory postural control, reactive postural control, sensory orientation, and dynamic gait.

### 3.3 Set shifting ability

The set shifting ability score was measured as the difference in time to complete Part B minus Part A of the Trail Making Test (Sanchez-Cubillo et al., 2009; McKay et al., 2018). Part A requires participants to quickly connect sequentially numbered dots (1-2-3, etc.) and is a test of visuomotor search and psychomotor speed. Part B is similarly formatted but requires participants to shift between numbers and letters (1-A-2-B, etc.), which tests the same domains with the additional requirement that participants shift between the numbers and letters. A greater difference in time to complete Part B compared to Part A indicates slower cognitive set shifting and therefore lower cognitive set shifting ability.

### 3.4 Overall cognitive ability

The Montreal Cognitive Assessment (MoCA, www.mocatest.org) was given as a rapid assessment of overall cognitive ability that assesses cognitive domains including executive function, attention, and memory (Nasreddine et al., 2005). Participants also self-reported the number of years of education. These data were collected as potential covariates so we could test whether overall cognitive ability confounds more specific associations to cognitive set shifting and were not considered for exclusion, as a range of cognitive abilities was necessary to achieve the goals of this study.

### 3.5 Perturbations

Participants were exposed to 48 translational support-surface perturbations of unpredictable timing, direction, and magnitude using a custom perturbation platform (Payne et al., 2019a). Perturbations were evenly divided between forward and backward directions, and three perturbation magnitudes, which will be referred to as small, medium, and large. The small perturbation (0.15 g, 11.1 cm/s, 5.1 cm) was identical across participants. Medium (0.21-0.22 g, 15.2-16.1 cm/s, 7.0-7.4 cm) and large (0.26-0.29 g, 19.1-21.0 cm/s, 8.9-9.8 cm) perturbations were linearly scaled down from reference magnitudes (medium: 0.22 g, 16.7 cm/s, 7.7 cm; large: 0.30 g, 21.8 cm/s, 10.2 cm) by multiplying perturbation acceleration, velocity, and displacement characteristics by a scaling factor linearly related to the participant’s height (Equation 1) to account for the effect of participant height and deliver perturbations that are mechanically similar across body sizes (Payne et al., 2019a). The 48 perturbation series was divided into 8 blocks, each containing one replicate of each unique perturbation. Three different block-randomized perturbation orders were used across participants to randomize any possible effects of trial order. The perturbations for an example participant are shown in Figure 1.

**Figure 1.**
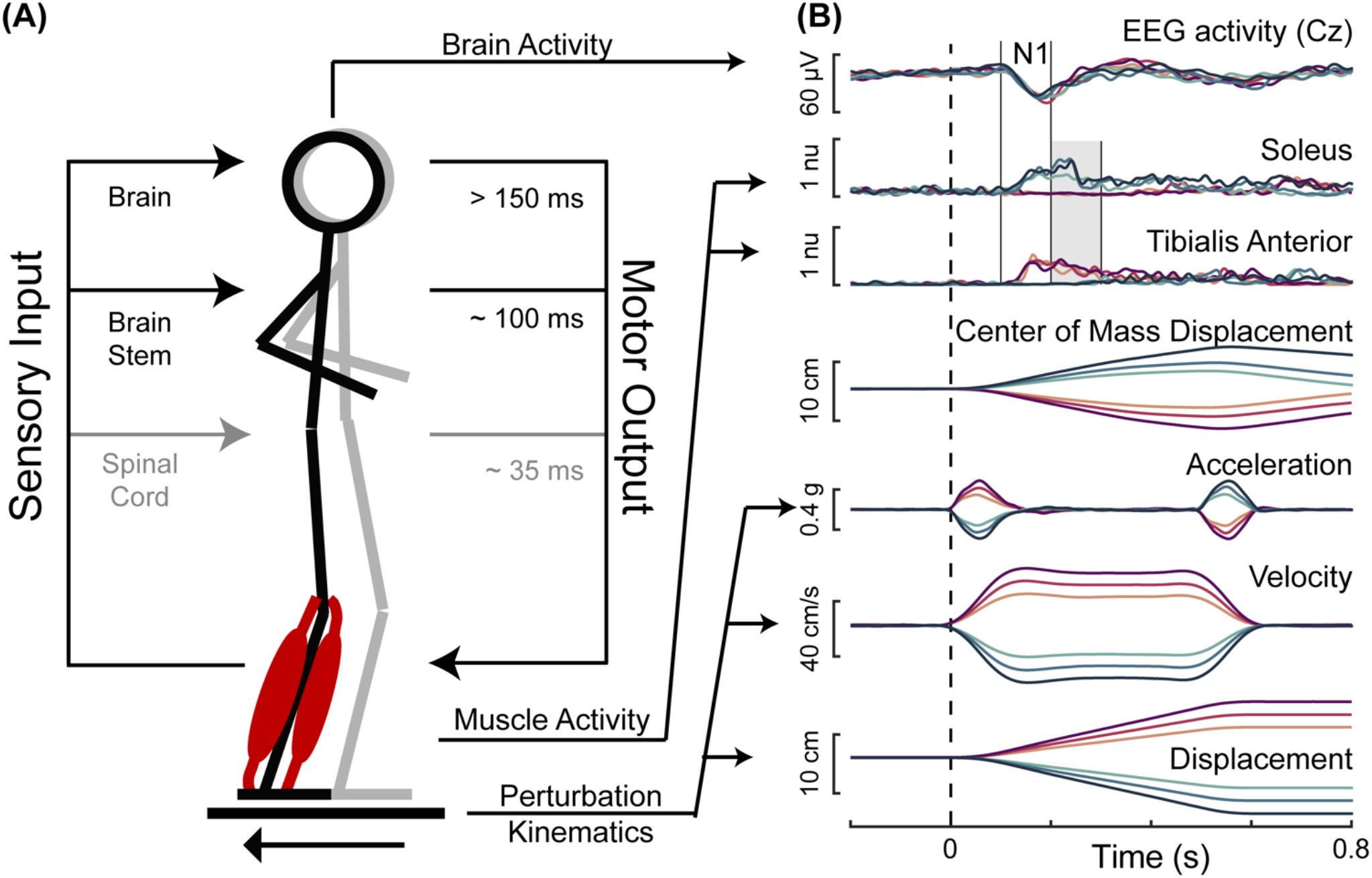
Balance perturbations. **(A)** The translational support-surface balance perturbation is depicted along with a schematic displaying hierarchical levels of control of the perturbation-evoked muscle activity. **(B)** Perturbation kinematics and the measured response variables are shown for an example participant with forward movements of the floor represented in magentas and backward movements of the floor represented in blues, with darker colors for larger perturbations. Perturbation onset is indicated with the dashed vertical line. Solid vertical lines indicate the time window of 100-200 ms, in which the cortical N1 and the early phase of muscle activity were assessed. The shaded gray area indicates the time window of 200-300 ms, in which the late phase of muscle activity was assessed.

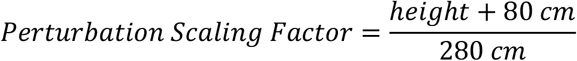

Participants were instructed to cross their arms across their chest, focus their vision on a fixed location at eye-level approximately 4.5 meters away, and to try to recover balance without taking a step. The experimenter monitored continuous electromyography (EMG) and electroencephalography (EEG) activity to ensure activity had returned to quiet baseline levels prior to the onset of the next perturbation. Seated rest breaks were taken after 15 minutes of perturbations, or more frequently at the request of the participant or if the participant displayed signs of fatigue or loss of concentration.

Excluding rest breaks, inter-trial-intervals, from perturbation onset to perturbation onset were 23 ± 11 s. Trials in which participants took steps (8% of all trials) were excluded from analysis.

### 3.6 Body motion

Motion of the body’s center of mass was tracked using a 10-camera Vicon Nexus 3D motion analysis system. Reflective markers placed on areas of the body including the head, neck, hips, knees, ankles, and feet were used to create a model of the body, and Vicon’s plug-in-gait model was used to estimate the body’s center of mass. Body motion was referenced to motion of the support-surface to assess the deviation of the center of mass from the base of support. Body motion was then quantified for each participant in each trial type as the peak deviation of the center of mass from the base of support along the axis of perturbation motion in data averaged across perturbations for each direction and magnitude.

### 3.7 Muscle activity

Surface EMGs (Motion Lab Systems, Baton Rouge, LA) were collected from tibialis anterior and soleus muscles, which are an agonist-antagonist pair of ankle muscles activated in forward and backward support-surface perturbations. EMG activity was collected using silver silver-chloride bipolar electrodes with 2 cm interelectrode spacing (Norotrode 20, Myotronics, Inc.) and sampled at 1000 Hz after an online analog 500 Hz low-pass filter. Skin was scrubbed with alcohol swabs and shaved if necessary prior to electrode placement.

Muscle activity was epoched into 2.4 second intervals, beginning 400 ms before perturbation onset. Epochs were 35 Hz high-pass filtered with a third-order zero-lag Butterworth filter, mean-subtracted, half-wave rectified, and then 40 Hz low-pass filtered. To avoid issues with normalization that could occur when averaging muscle activity across the left and right legs, only the muscle activity for the left leg was analyzed. As only nonstepping behaviors were analyzed in forward and backward perturbations, muscle activity is expected to be similar across left and right legs.

Muscle activity was assessed in two time bins of interest. This included an early (100-200 ms) time bin, which is expected to primarily contain involuntary brainstem-mediated activity, and a late (200-300 ms) time bin, in which cortical contributions to muscle activation can appear (Figure 1).

Antagonist muscle activation was quantified relative to agonist muscle activity (Lang et al., 2019) within each perturbation magnitude as a measure of directional specificity, or motor set shifting, in terms of how the same muscle is activated differently across perturbation directions. EMG activity for each muscle was averaged within each time bin for each of the perturbation magnitudes and directions. Then, EMG activity was quantified according to Equation 2, which takes the absolute value of the difference in EMG activity between forward and backward perturbation directions (i.e., agonist activity - antagonist activity) and divides it by the larger of the two values (i.e., agonist activity). This results in a value between 0 and 1, where values near 1 indicate nearly exclusive agonist activity, or high directional specificity, and values near zero indicate nearly identical agonist and antagonist activity, or low directional specificity.

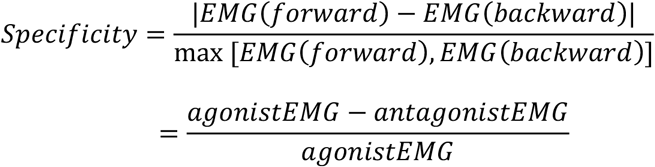

### 3.8 Cortical activity

Electroencephalography (EEG) data were collected during the perturbation series using thirty-two active electrodes (ActiCAP, Brain Products, Germany) placed according to the international 10-20 system, with the exception of two electrodes placed on the skin over the mastoid bones behind the ears for offline re-referencing. The electrodes were prepared with conductive electrode gel (SuperVisc 100 gr. HighViscosity Electrolyte-Gel for active electrodes, Brain Products) using a blunt-tipped needle, which was also used to rub the scalp to improve signal quality. Impedances for the Cz and mastoid electrodes were generally below 10 kOhm before the start of data collection.

To subtract vertical eye movement and blink artifacts, electrooculography (EOG) data were collected with bipolar passive electrodes (E220x, brain Products) vertically bisecting the right pupil and referenced to the forehead. Prior to placement, the EOG electrodes were prepared with high-chloride abrasive gel (ABRALYT HiCl 250 gr., High-chloride-10% abrasive electrolyte gel, Brain Products) and the skin was scrubbed with an alcohol swab. EEG and EOG data were sampled at 1000 Hz on an ActiCHamp amplifier (Brain Products) with a 24-bit A/D converter and an online 20 kHz anti-aliasing low-pass filter.

EEG data were 1 Hz high-pass filtered offline with a third-order zero-lag Butterworth filter, mean-subtracted within channels, and then low-pass filtered at 25 Hz. Data at the Cz electrode were re-referenced to the average of the two mastoid electrodes and epoched into 2.4 s segments beginning 400 ms before perturbation onset. EOG data were similarly filtered and segmented without re-referencing. The Gratton and Coles (Gratton et al., 1983) algorithm was applied as described in Payne et al. (Payne et al., 2019a) to remove blinks and eye movement artifacts through a serial regression-subtraction approach. Cz epochs were then averaged across trials within each trial type and baseline subtracted using a baseline period of 50-150 ms before perturbation onset. Cortical N1 response amplitudes were then quantified as the negative of the most negative amplitude (µV, such that higher values indicate a larger component peak) at the Cz electrode between 100-200 ms after perturbation onset (Figure 1).

### 3.9 Statistical analyses

Simple linear regressions were used to test for associations between pairs of study variables. Specifically, linear regressions were used to test set shifting ability scores as a predictor of: N1 amplitudes, peak center of mass displacements, and directional specificity of soleus and tibialis anterior muscle activity. Additional linear regressions were used to test each of these variables as a predictor of MiniBESTest scores. Variables that were not normally distributed as determined by Shapiro-Wilk test p-values<0.05 were transformed to a normal distribution prior to regression using boxcox.m in MATLAB. Parameter estimates for the regression slopes were compared against the hypothesized value 0 with two-sided t-tests using PROC GLM in SAS. Figures display untransformed data with p-values and R^2^ values obtained from the adjusted variables.

All tests for association with N1 amplitudes and peak center of mass motion were performed separately across the two perturbation directions and three perturbation magnitudes, and tests for association with EMG activity were performed separately across the three perturbation magnitudes. Associations were examined for consistency across testing conditions via visual inspection of regression plots and tabulated regression coefficients. Simple linear regressions that yielded significant associations were further tested for robustness to potential confounding variables, including age, sex, height, weight, overall cognition scores, years of education, and balance ability scores. Specifically, each significant regression was retested in a multivariate regression with each of the potential confounding variables added, one at a time, as an additional predictor in the model to confirm that the associations were insensitive to adjustment by the potential confound.

## 4. RESULTS

Demographic characteristics of the participants are shown in Table 1. Overall, participants had high clinical balance ability and cognitive set shifting ability. On the clinical balance test, most participants scored above the fall-risk threshold of 23 for adults between 70-80 years old (Magnani et al., 2020) (Figure 2). Similarly, most participants performed better on the Trail Making Test (B-A) than the average of 46 s that would be expected for adults between 70-80 years old with 12+ years of education (Tombaugh, 2004). Clinical balance ability scores were not associated with any other study variable. Specifically, miniBESTest scores were not associated with cognitive set shifting (p=0.26, Figure 2), the peak amplitude of center of mass displacement (p>0.12 across all perturbation magnitudes directions), antagonist activity of the soleus (all p>0.078), or tibialis anterior (all p>0.50), or the peak amplitude of the cortical N1 response (all p>0.57).

**Table 1.** Participant characteristics.

|  |  |
| --- | --- |
| N | 19 |
| Sex | 6F, 13M |
| Age | $71 \pm 7$ |
| Height (cm) | $175 \pm 10$ |
| Weight (kg) | $79 \pm 16$ |
| Trail Making Test (part A, seconds) | $24 \pm 8$ |
| Trail Making Test (part B, s) | $61 \pm 25$ |
| Set Shifting (part B-A, s) | $37 \pm 20$ |
| Overall Cognition (MoCA, /30) | $26 \pm 3$ |
| Years of Education | $17 \pm 2$ |
| Balance Ability (miniBESTest, /28) | $25 \pm 2$ |

**Figure 2.**
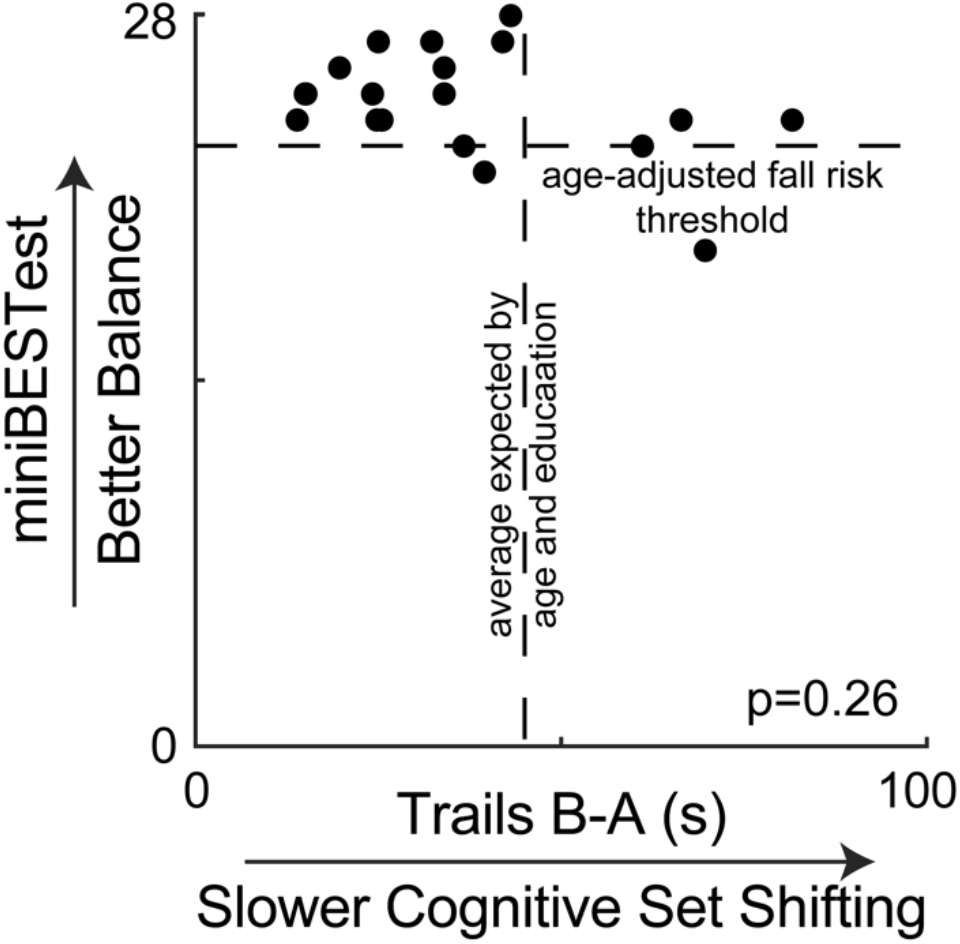
Clinical balance scores are plotted against cognitive set shifting performance, showing no association. Most participants scored above the suggested fall risk cutoff score of 23 for people between 70-80 years old on the miniBESTest (Magnani et al., 2020). Most participants also completed the Trail Making Test (B-A) faster than the average of 46 seconds that would be expected by their age and education from normative data (Tombaugh, 2004).

Lower cognitive set shifting ability was associated with stiffer responses to perturbations (Figure 3). Specifically, individuals who took longer to complete the cognitive set shifting task had smaller peak amplitudes of center of mass displacement with respect to the base of support in all perturbation magnitudes in both perturbation directions (forward perturbations: small p=0.002 R^2^=0.44, medium p=0.001 R^2^=0.48, large p=0.003 R^2^=0.42; backward perturbations: small p<0.001 R^2^=0.49, medium p<0.001 R^2^=0.52, large p<0.001 R^2^=0.66). Set shifting ability scores remained a significant predictor of center of mass displacement in all perturbation magnitudes and directions when potential confounding variables of age, sex, height, weight, overall cognition, education, and balance ability scores were included in the models (Supplemental).

**Figure 3.**
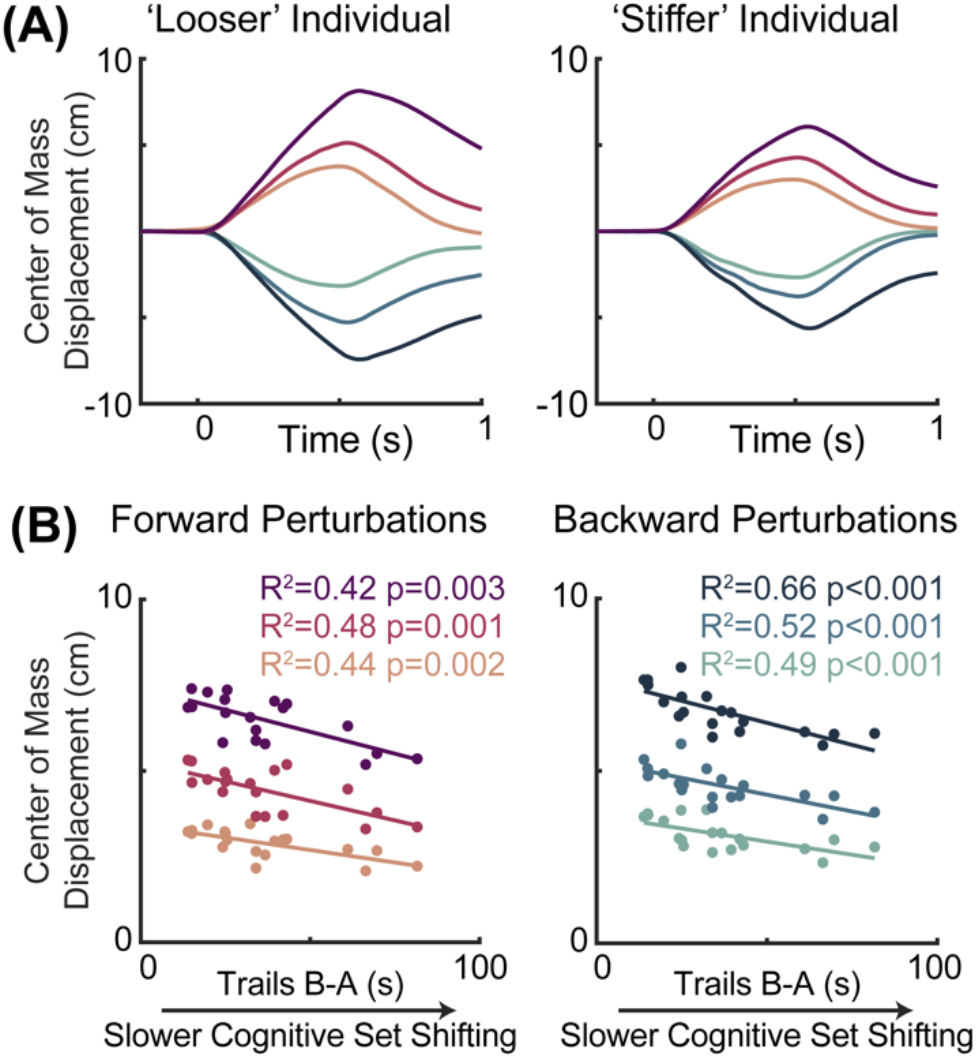
Slower cognitive set shifting was associated with stiffer behavioral responses to perturbation. **(A)** Center of mass displacements are shown for a looser individual (left) and a stiffer individual (right) for each of the perturbation types, with forward movements of the floor (resulting in backward leaning posture) represented in magentas and backward movements of the floor represented in blues, with darker colors for larger perturbations. **(B)** Peak center of mass displacements are plotted against cognitive set shifting scores for forward (left) and backward (right) perturbations for each perturbation magnitude.

Lower cognitive set shifting ability was associated with more antagonist activity in the late phase (200-300 ms) of soleus muscle activation (Figure 4). Specifically, individuals who took longer to complete the cognitive set shifting task displayed less directional specificity of late phase (200-300 ms) soleus activity (small perturbation p=0.010, R^2^=0.33, medium p=0.007, R^2^=0.35, large p=0.004 R^2^=0.39). Set shifting ability scores remained a significant predictor of directional specificity of late phase soleus activity in all perturbation magnitudes when potential confounding variables of age, sex, height, weight, overall cognition, education, and balance ability scores were included in the models (Supplemental). In contrast, set shifting ability was not associated with directional specificity of the early automatic phase (100-200 ms) of soleus muscle activation (small perturbation p=0.12, medium p=0.41, large p=0.16).

**Figure 4.**
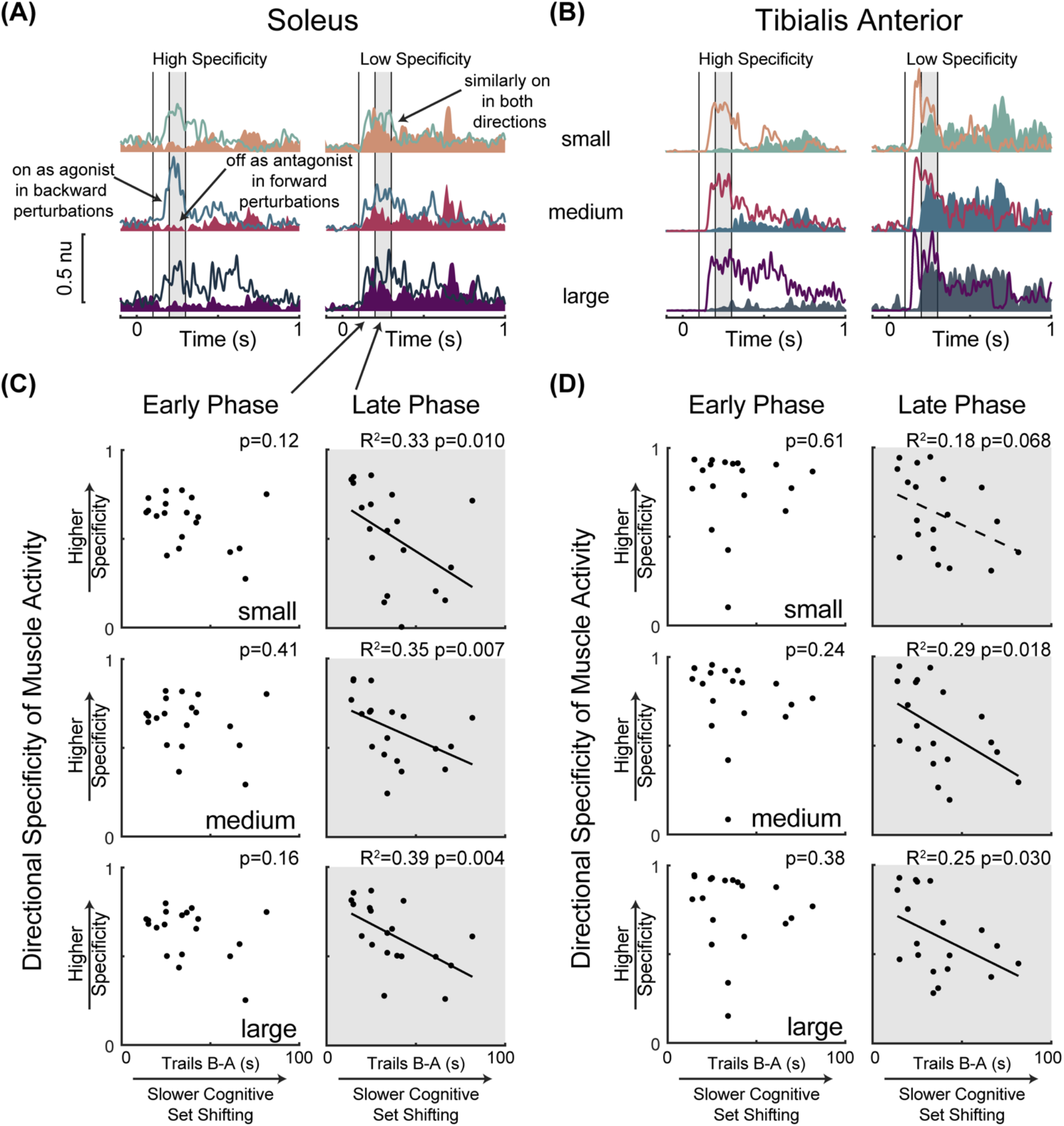
Slower cognitive set shifting was associated with lower directional specificity in the late phase of muscle activation. Condition-averaged muscle activity is shown for **(A)** soleus and **(B)** tibialis anterior muscles for example participants with higher or lower directional specificity scores. Muscle activity evoked by forward movements of the floor (resulting in backward leaning posture) is represented in magentas and activity evoked by backward movements of the floor is represented in blues, with darker colors for larger perturbations. Antagonist activity (i.e., soleus activity in forward perturbations and tibialis anterior activity in backward perturbations) is shaded for clarity. Vertical lines at 100 ms, 200 ms, and 300 ms mark the bounds of the time bins of interest. The later (200-300 ms) time bin is shaded in all panels. Directional specificity is plotted against cognitive set shifting scores for the **(C)** soleus muscle and **(D)** tibialis anterior muscle in each perturbation magnitude for early (100-200 ms) and late (200-300 ms) time bins.

Limited associations between cognitive set shifting ability and tibialis anterior muscle activity (Figure 4) were not robust to the inclusion of potential confounding variables. Specifically, individuals who took longer to complete the cognitive set shifting task displayed lower directional specificity of late phase (200-300 ms) tibialis anterior muscle activation in medium (p=0.018 R^2^=0.29) and large (p=0.030 R^2^=0.25) perturbations, but only a trend was observed in small (p=0.068) perturbations. However, these associations were not robust to the inclusion of potential confounding variables (Supplemental). Specifically, set shifting scores were no longer a significant predictor of directional specificity of late phase tibialis anterior muscle activation in the medium perturbation magnitude upon the inclusion of sex (p=0.053) or overall cognition (p=0.096) into the model, and significance was similarly lost in the large perturbation magnitude upon the inclusion of height (p=0.058), sex (p=0.11), or overall cognition (p=0.077) into the model. Set shifting ability scores were not significant predictors of directional specificity of the early automatic phase (100-200 ms) of tibialis anterior muscle activation (small perturbation p=0.61, medium p=0.24, large p=0.38).

Lower cognitive set shifting ability was associated with larger perturbation evoked cortical N1 responses (Figure 5). Specifically, individuals who took longer to complete the cognitive set shifting task had larger cortical N1 peak amplitudes in response to all perturbation magnitudes in both perturbation directions (forward perturbations: small p=0.004 R^2^=0.40, medium p=0.002 R^2^=0.44, large p=0.003 R^2^=0.41; backward perturbations: small p<0.001 R^2^=0.60, medium p=0.004 R^2^=0.39, large p=0.016 R^2^=0.30). Set shifting ability scores remained a significant predictor of N1 peak amplitudes in all perturbation magnitudes and directions when potential confounding variables of age, sex, height, weight, overall cognition, education, and balance ability scores were included in the models, with one exception (Supplemental). This exception was in large backward perturbations, where set shifting ability fell below significance as a predictor of N1 amplitudes upon inclusion of sex into the model (p=0.055).

**Figure 5.**
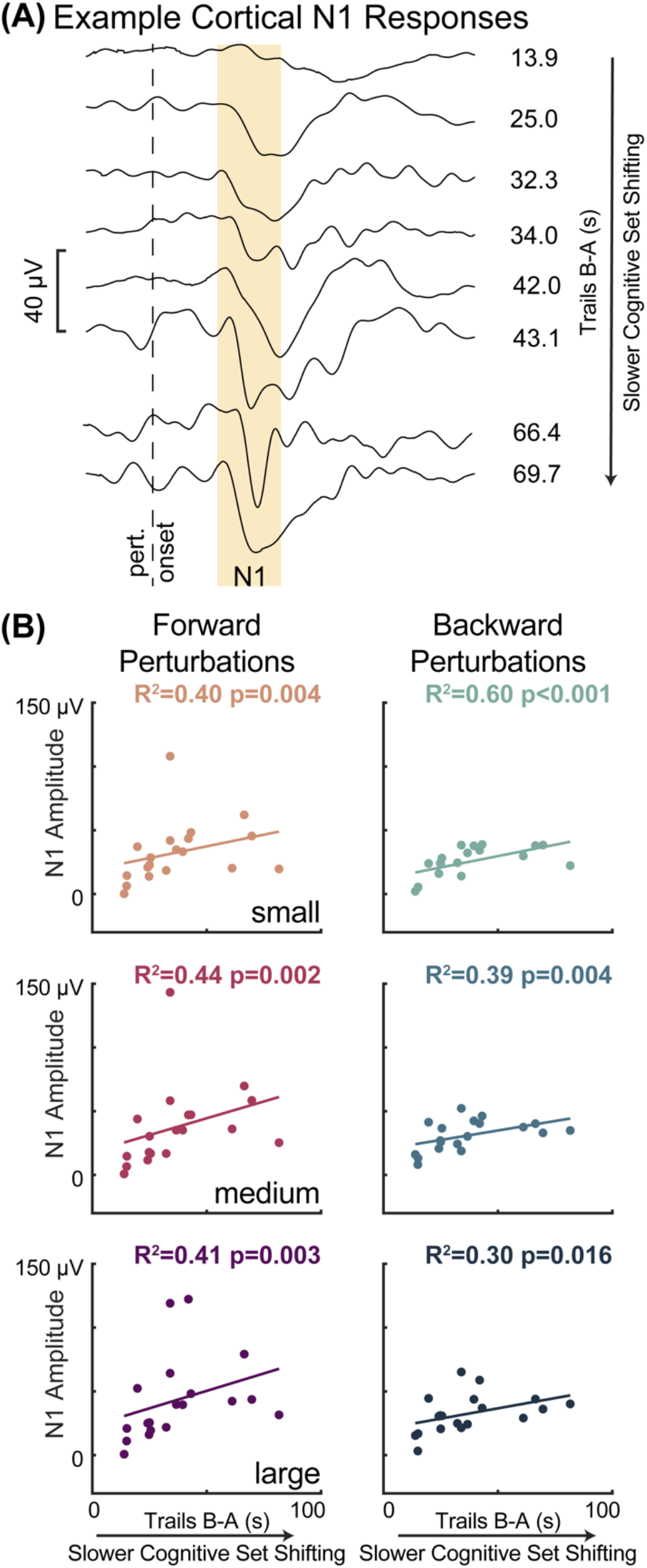
Slower cognitive set shifting was associated with larger perturbation-evoked cortical N1 responses. **(A)** Cortical responses are shown at the Cz electrode for eight different participants (averages across all trials) as examples along with their set shifting scores. The gold box highlights the window of 100-200 ms, in which the N1 amplitude was quantified. **(B)** Cortical N1 response amplitudes are plotted against cognitive set shifting scores in each of the perturbation conditions. Forward movements of the floor (resulting in backward leaning posture) are represented in magentas and backward movements of the floor are represented in blues, with darker colors for larger perturbations.

## 5. DISCUSSION

Our results suggest that cortically-mediated reactions to a balance disturbance may share common mechanisms with cognitive set shifting ability in older adults with relatively high balance ability. While prior studies have linked set shifting impairments to severe balance dysfunction and frequent falls (Herman et al., 2010; Tangen et al., 2014; McKay et al., 2018), our results demonstrate that set shifting ability is expressed in successful balance recovery behavior even in the absence of profound clinical balance disability. Individuals with lower cognitive set shifting ability had stiffer whole-body behavior after balance perturbations, which may be caused by excessive agonist-antagonist coactivation related to difficulty incorporating the directional context into the cortically-mediated phase of the motor response. The associations between cognitive set shifting and the late phase of the motor response as well as the larger cortical N1 responses both suggest that cognitive set shifting shares common mechanisms with the cortically-mediated reaction to the perturbation, rather than cortical preparatory activity, which would have been observed in the early phase of the motor response. The associations between balance and cognitive set shifting cannot be explained by age, as all associations remained significant when accounting for age. Our data suggest that cognitive set shifting ability may indicate variation in balance control that is not yet detectable in the clinical balance test, but it remains to be tested whether cognitive set shifting ability could serve as an earlier predictor of falls in preclinical populations. Earlier detection of changes in balance control could enable treatment when these changes are more amenable to adaptation (Zaback et al., 2019). Further, new therapeutic targets for rehabilitation could be identified if there are causal links between the engagement of cortical resources and the subsequent antagonist muscle activity, which would need to be tested with methods that disrupt the cortical activity to influence balance and cognitive set shifting behavior. These findings warrant further investigation of cortical engagement in balance recovery behavior guided by cognitive set shifting ability rather than just clinical balance ability, which may help explain why balance rehabilitation can be enhanced by cognitive training (Hagovska and Olekszyova, 2016).

These findings suggest that cognitive set shifting ability is expressed in balance recovery behavior earlier than previously suggested. The clinical balance test (Balance Evaluation Systems Test, or BESTest, later shortened to the miniBESTest) was designed to distinguish between levels of balance function and fall risk in people seeking treatment for balance-related disability (Horak et al., 2009; Franchignoni et al., 2010). The miniBESTest was not designed to measure subtle differences in balance recovery behavior in preclinical balance function, which may explain why we did not find associations between clinical balance ability and any other variable in the present study. By assessing more precise neuromechanical metrics of balance control in response to balance destabilization, we found that cognitive set shifting ability was associated with stiffer balance recovery mechanics and antagonist muscle activity that were not observed in the clinical balance test. While the majority of our participants had a high level of balance function, we chose not to exclude people with mild cognitive impairment to allow for variation in cognitive function to relate to balance recovery behavior. Although our cognitive set shifting scores were typical given the age and relatively high level of education of our participant cohort (Giovagnoli et al., 1996; Tombaugh, 2004; Steinberg et al., 2005), this should not be misinterpreted as an indication that our participants are free of the cognitive decline that would be expected for their ages. Indeed, roughly a quarter of our participants (5 of 19) scored below the cutoff value for mild cognitive impairment on the Montreal Cognitive Assessment (Nasreddine et al., 2005). In any case, most of the observed associations between cognitive set shifting and perturbation-evoked balance recovery behavior remained significant when accounting for age, education, overall cognition, and most of the other potentially confounding variables considered, indicating that our associations are specific to cognitive set shifting and not better explained by these other factors.

Difficulty shifting cognitive sets may extend to a related difficulty shifting motor sets in the cortically-mediated phase of balance recovery behavior. People with lower cognitive set shifting ability had stiffer whole-body mechanics as evidenced by less center of mass motion in response to balance perturbations. While greater resistance to a balance disturbance would seem to suggest greater stability (Horak et al., 2005), the accompanying increase in antagonist activity, which is associated with clinical balance impairments (Lang et al., 2019), suggests this biomechanical stiffness does not reflect better balance control. Our directional specificity measure of antagonist activity was originally developed as a proxy measure for simultaneous cocontraction of agonist-antagonist muscle pairs that overcomes issues of comparing activity levels between muscles by instead comparing the activity of an individual muscle between agonist and antagonist contexts (Lang et al., 2019). However, because this measure compares the shift in activity of individual muscles between agonist and antagonist contexts, it could also be considered as a measure of motor set shifting. Directional specificity of the late phase of the soleus muscle was robustly associated with cognitive set shifting ability across perturbation magnitudes and robust to all potential confounds, but associations with the late phase of the tibialis anterior muscle were limited to larger perturbations and may be explained by lower overall cognition or female sex. While these muscles are a small fraction of the muscle activity contributing to the overall balance recovery behavior, our data provide evidence that difficulty shifting cognitive sets may extend to a related difficulty shifting motor sets between perturbation directions in the cortically-mediated phase of the motor response. A relationship between cognitive and motor set shifting is further supported by a recent finding that older adults with difficulty shifting cognitive sets had a related difficulty shifting between locomotor patterns on a split belt treadmill (Sombric and Torres-Oviedo, 2021). Because the motor set shifting association was not observed in the early phase of the perturbation-evoked motor response, it is unlikely that the influence of pre-perturbation cognitive state, such as anticipation, readiness, or arousal, on the brainstem-mediated response has overlapping mechanisms with cognitive set shifting. The association of motor set shifting during the later cortically-mediated phase of the motor response suggests that instead the cortically-mediated reaction of the perturbation may have overlapping mechanisms with cognitive set shifting, which is further supported by the enhanced cortical N1 responses.

Our findings suggest the cortical N1 has overlapping mechanisms with cognitive set shifting, and prior work linking the N1 to attention and perceived threat may help explain the subsequent antagonist activity. While prior studies have shown within-subjects changes in the cortical N1 with cognitive processes including attention (Quant et al., 2004b; Little and Woollacott, 2015), perceived threat (Adkin et al., 2008; Mochizuki et al., 2010), and predictability (Adkin et al., 2006; Adkin et al., 2008; Mochizuki et al., 2008; Mochizuki et al., 2010), we believe this is the first study to demonstrate an association to between-subjects differences in cognitive ability. Given that set shifting ability is reflected in the cortical activity (100-200 ms) prior to the muscle activity (200-300 ms), it is possible that the N1 reflects cortical activity contributing to the subsequent antagonist activity, which could be tested in future studies by modulating cortical activity through therapeutic noninvasive brain stimulation (Taube et al., 2006; Jacobs et al., 2009). For example, after the perturbation, the participant may perceive a high threat or need for attention, which is reflected in the cortical N1 (Quant et al., 2004b; Adkin et al., 2008; Mochizuki et al., 2010; Little and Woollacott, 2015), and subsequently engage a nonspecific cocontraction strategy that does not incorporate the directional context into the behavior. Indeed, people report paying more attention to balance under more threatening conditions (Huffman et al., 2009) and display greater stiffness of postural sway (Carpenter et al., 2001; Carpenter et al., 2004; Carpenter et al., 2006). Greater perceived threat or fear of falling is also associated with agonist-antagonist cocontraction in both younger and older adults (Okada et al., 2001; Carpenter et al., 2006). Further, habituation of agonist-antagonist cocontraction with practice in high threat conditions is associated with habituation of the emotional response (Zaback et al., 2019), which may be easier to modify than the abnormal involuntary behavior observed at more severe stages of balance impairment (McKay et al., 2021). We previously suggested that the cortical N1 could reflect compensatory cortical control based on larger amplitudes in young adults with lower balance ability (Payne and Ting, 2020a) and on trials with compensatory steps (Payne and Ting, 2020c). This nonspecific cocontraction could be another way in which compensatory cortical control is engaged. Although the supplementary motor area has direct connections to motor neurons (Goldberg, 1985), potential connections between the cortical N1 and subsequent muscle activation also include indirect routes, such as through projections from the supplementary motor area to the motor cortex or basal ganglia (Goldberg, 1985). However, any causal links between the cortical N1 and balance recovery behavior remain speculative until tested by further studies, particularly through methods that would disrupt the cortical activity, such as noninvasive brain stimulation, or dual task interference, which reduces the N1 amplitude (Quant et al., 2004b; Little and Woollacott, 2015) and the late phase of the muscle activity (Rankin et al., 2000).

Changes in prefrontal cortical activity in older adults may explain links between motor and cognitive behavior and may be a potential target for rehabilitation. Cognitive set shifting depends on the dorsolateral prefrontal cortex (Zgaljardic et al., 2006), and the cortical N1 has been localized to the supplementary motor area (Marlin et al., 2014; Mierau et al., 2015), but there are several potential explanations as to why cognitive set shifting would be associated with the cortical N1 response despite their distinct brain regions. First, older adults recruit prefrontal cortical areas to a greater extent and more broadly than young adults for the same tasks (Reuter-Lorenz and Cappell, 2008), and lose functional segregation between different cortical areas (Chen et al., 2011; Damoiseaux, 2017; Chong et al., 2019; Cassady et al., 2020), which may result in coupled activation between cognitive and motor cortical areas. Accordingly, older adults tend to recruit prefrontal cortical areas broadly for balance and walking tasks (Stuart et al., 2018; Nobrega-Sousa et al., 2020; St George et al., 2021). However, the cortical N1 may not arise exclusively from the supplementary motor area, as there is evidence to suggest that multiple cortical sources synchronize in the theta frequency band to contribute to the cortical N1 response even in young adults (Peterson and Ferris, 2018; 2019). Increased synchronization between prefrontal and motor cortical areas during balance recovery with aging may explain associations between cognitive function and balance control in older adults. For example, we recently showed that beta coherence between motor and prefrontal cortical areas during balance recovery in older adults is associated with cognitive dual task interference in walking (Palmer et al., 2021). A better understanding of the mechanisms linking balance and cognitive function in aging could reveal new therapeutic targets for rehabilitation and enable a more targeted exploration of the effects of cognitive training on balance rehabilitation (Smith-Ray et al., 2015; Hagovska and Olekszyova, 2016). For instance, it is well established that noninvasive stimulation of the dorsolateral prefrontal cortex can affect cognitive set shifting performance (Ko et al., 2008a; Ko et al., 2008b; Leite et al., 2011; Leite et al., 2013; Luthi et al., 2014; Gerrits et al., 2015; Tayeb and Lavidor, 2016; Imburgio and Orr, 2018; Leite et al., 2020), but similar stimulation protocols are rarely applied to impact balance function despite evidence that it can reduce cognitive dual task interference on balance and walking behaviors (Manor et al., 2018).

## Respective Contributions

A.M.P. and L.H.T. conceived and designed the experiment, A.M.P. collected the data, performed all analyses, drafted and revised the manuscript and figures, J.A.P. and J.L.M. contributed to the data analysis, L.H.T. and J.A.P. contributed to the interpretation of results and manuscript revision, and all authors approved of the final manuscript.

